# Simultaneous determination of ^87^Sr/^86^Sr and trace-element data in otoliths and other sclerochronological hard structures

**DOI:** 10.1101/2020.04.24.060640

**Authors:** Jens C. Hegg, Christopher M. Fisher, Jeffrey Vervoort

**Affiliations:** Department of Fish & Wildlife Sciences, University of Idaho, Moscow, ID 83844-1136, USA; School of the Environment, PO Box 642812, Washington State University, Pullman WA 99164-2812, USA; Centre for Exploration Targeting, School of the Earth Sciences, The University of Western Australia, Crawley, WA 6009, Australia

**Keywords:** sclerochronology, trace elements, strontium ratio, carbonate, otolith, inductively coupled plasma mass-spectrometry, LASS analysis

## Abstract

Chronological data from hard structures have been instrumental in reconstructing information about the past across numerous disciplines. Isotopic and trace elemental chronologies from the depositional layers of speleothems, corals, bivalve shells, fish otoliths and other structures are routinely used to reconstruct climate, growth, temperature, geological, archeological and migratory histories. Recent *in situ* analytical advances have revolutionized the use of these structures. This is particularly true of fish, in which detailed origin, life-history, and migration history can be reconstructed from their otoliths. Specifically, improvements in laser ablation-inductively coupled plasma mass spectrometry (LA-ICPMS) have allowed increases in temporal resolution, precision, and sample throughput. Many studies now combine multiple chemical and isotopic tracers, taking advantage of multivariate statistical methods and multiple trace-elements and isotope systems to glean further information from individual samples. This paper describes a novel laser ablation split-stream (LASS) methodology which allows simultaneous collection of the Sr isotope composition (^87^Sr/^86^Sr) and trace-elemental data from chronologically deposited carbonate samples. The study investigates the accuracy and precision of varying laser spot sizes on a marine shell standard and fish otoliths using LASS and presents a comparison to traditional “single stream methods” using pre-existing otolith data on the same samples. Our results indicate that LASS techniques can be used to provide accurate and precise data at the same laser spot sizes as previous otolith studies, thereby doubling analytical throughput, while also providing improved spatially and temporally-matched data reduction using newly developed features for the Iolite data reduction platform.

## Introduction

Chronological data from hard structures have been instrumental in reconstructing information about the past across numerous disciplines. While tree ring dendrochronology is one of the most well-known, isotopic and trace elemental chronologies from the depositional layers of speleothems have been instrumental in reconstructing climate and temperature records from the near and deep past (Hendy and Wilson 1968, Fairchild et al. 2012). Geochemistry of dust trapped in ice cores has been used to reconstruct global wind patterns (Rodda et al. 2019). Chemical and structural sclerochronologies from bivalve shells, corals, and other animal hard parts are used to reconstruct climate, migration, and animal food sources with broad application to biology, archeology and climate science (Hudson et al. 1976, Oschmann 2009, Andrus 2011).

Understanding the provenance and movement patterns of migratory organisms is often critical to understanding their ecology (Hobson 1999). Increasingly, hard-parts from the animals themselves have been found to store chemical records of these movements (Wassenaar and Hobson 1998, Rubenstein and Hobson 2004, Hobson et al. 2010). These endogenous records can provide important information about the origin and migratory path of an individual, information that is often difficult, or impossible, to obtain through traditional non-in situ methods (Hobson et al. 2010). From birds to insects and mammals, endogenous chemical records of migration have revolutionized our understanding of animal movement (Webster et al. 2002). These chemical and isotopic sclerochronological approaches are now routinely used to reconstruct important aspects of biology and ecology including movement, ontogeny, population ecology and archeological patterns of human interaction with organisms (Jacques et al. 2002, Thornton 2011, Kohn et al. 2013, Hegg et al. 2015).

Endogenous chemical records have been particularly revolutionary in our understanding of fish movements and provenance (Begg et al. 2005, Secor 2010, Morales-Nin and Geffen 2015). The development and methods in this field mirror similar advances in related sclerochronological and speleothem fields. Fish otoliths grow in sequential layers of biogenic aragonite, preserving trace elements and isotopes unique to the environments the fish inhabits through its life (Campana 1999, Campana and Thorrold 2001). These chemical tracers have been used to uncover the population structure of fish populations (Walther and Thorrold 2008, Muhlfeld et al. 2012, Brennan and Schindler 2017), as well as the details of their migratory paths (Hegg et al. 2015, 2019, Sturrock et al. 2015b, Chittaro et al. 2019), sometimes with incredible precision on the landscape (Hamann and Kennedy 2012). Otoliths thus provide a high-resolution, broad scale, dataset of location, growth, and ontogeny across the life of the organism which is not possible in using tagging or tracking technologies (Hegg et al. 2018).

The increasing detail and resolution of information obtained from otoliths has been tied in large part to technological advances in laser ablation-inductively coupled plasma mass spectrometry (LA-ICPMS). Technological advances in instrumentation and sample chamber design have increased the resolution and throughput (Pozebon et al. 2017, Lobo et al. 2018). Analytical precision and signal-to-noise ratios have steadily improved in new ICPMS models, and new laser ablation systems have increased sampling efficiency, capacity and sample imaging resolution. Despite these advances, few publications referencing methodological improvements in otolith studies have been published since Barnett-Johnson et al. (2005) validated LA-ICPMS methods in otoliths.

These technological improvements have spurred important advances in otolith research and the percentage of papers using LA-ICPMS technology has markedly increased over the last two decades (Campana and Thorrold 2001, Begg et al. 2005, Miller et al. 2010, Walther 2019, Wang et al. 2019). Early studies relied on differences on dissolution of whole otolith samples for trace element or isotopic chemistry, or on relatively time and labor-intensive techniques to micro-mill material from individual otoliths bands (Campana et al. 1997, Volk et al. 2000, Kennedy et al. 2002). The introduction of laser ablation-inductively coupled plasma mass spectrometry (LA-ICPMS) allowed researchers to sample otoliths with greater spatial resolution—and with drastically higher throughput—greatly increasing the knowledge gained from otoliths and allowing more robust statistical interpretation due to markedly increased sample sizes (Outridge et al. 2002, Barnett-Johnson et al. 2005, Ludsin et al. 2006).

Currently, otolith studies using LA-ICPMS follow two analytical approaches. The first is analysis of trace elemental concentrations relative to calcium (expressed as a ratio) in the otolith matrix. Changes in these elemental ratios derive from the chemistry of the environment and ontological changes (Bath et al. 2000, Sturrock et al. 2015a). These unique ratios can then be used to reconstruct the location and conditions fish experience through their life (Walther et al. 2008). Otolith elemental ratios are most often measured on a sector field or quadrupole LA-ICPMS system, but other backscatter methods are also used (Campana et al. 1997, Limburg et al. 2013). A second, and complimentary, approach is the measurement of ^87^Sr to ^86^Sr in the otolith (^87^Sr/^86^Sr), which can be used as a high-resolution tracer of location for fish in fresh water (Kennedy et al. 1997). Strontium isotope compositions require measurement on a multicollector LA-ICPMS (Outridge et al. 2002, Barnett-Johnson et al. 2005, Woodhead et al. 2005).

Increasingly, researchers across the fields of animal hard-part and speleothem chronologies are using the improved analytical precision to understand chronologies with increasing resolution (Warter and Müller 2017). Scientists are also applying novel isotopic systems to reconstruct chronologies which in the past were confined largely to trace elements (Rüggeberg et al. 2008, Füllenbach et al. 2015), and combining isotopic compositions and trace element approaches to uncover multiple temporal records within the same structure (Zhou et al. 2009, Griffiths et al. 2010, Mallela et al. 2011, Woodcock et al. 2013, Chittaro et al. 2019).

A larger suite of analytes is often useful to reconstruct greater chronological detail. For this reason the use of multicollector ICPMS in combination with sector field or quadrupole ICPMS is increasingly employed in otolith studies (Zitek et al. 2010, Martin et al. 2013, Crook et al. 2013, Garcez et al. 2014, Hegg et al. 2018, 2019). This multivariate approach, while analytically powerful, increases both the cost and the time required for analysis since each otolith must be ablated twice using the traditional ‘single stream’ (SS) technique. This SS technique requires collecting ^87^Sr/^86^Sr on an MC-ICPMS instrument and trace-element data on a second quadrupole or sector-field ICPMS using a second laser transect. Further, we have found significant problems can arise in matching the two data-streams, since each analysis requires its own laser path on the otolith, and the two techniques typically require differing integration times. Controlling for these methodological mismatches can further increase the time and complexity involved in combining trace-element and ^87^Sr/^86^Sr analyses.

These alignment problems have been solved in other geochemical applications using laser ablation split-stream (LASS) methods to simultaneously collect data from one sample on two instruments (Fisher et al. 2014, Goudie et al. 2014). LASS has also been used in a single study in otoliths to measure ^87^Sr/^86^Sr and trace-elements with a notably large 150 μm laser spot diameter and using manual alignment of data (Prohaska et al. 2016).

The current study presents a LASS methodology for collection and data reduction of ^87^Sr/^86^Sr and multiple trace elements across otolith transects resulting in data whose integrations are matched both spatially and temporally on the sample. The methods are generally applicable to any carbonate sample requiring the combined measurement of trace elements and ^87^Sr/^86^Sr, particularly to sclerochronological and speleothem analysis. Our first objective is to present a novel laser ablation split-stream (LASS) method for simultaneous collection of ^87^Sr/^86^Sr and trace-element data. Second, we demonstrate the analytical precision of this method using the latest generation of MC-LA-ICPMS. To do so we compare the results of increasingly smaller laser spot sizes on two standard materials. Further, we compare the ‘real world’ precision of otolith LASS and SS ^87^Sr/^86^Sr analysis with prior results through a re-analysis of otolith samples analyzed in a prior study on older equipment in single stream configuration (Hegg et al. 2018).

## Methods

### Experimental Design

To test the accuracy and precision of the LASS method, results were compared from analysis of a homogeneous modern marine shell standard (MMS) material as well as otolith material, across a range of laser spot sizes. In a first experiment, the MSS was ablated sequentially using parallel lines comprising 100 seconds of data in each ablation, at laser spot diameters of 40, 35, 25, 20, 15, and 10 μm using both LASS and SS methods.

In a second experiment, four otoliths of adult fall Chinook salmon (*Oncorhynchus tshawytscha*) prepared at the Kennedy LIFE Lab at University of Idaho from a prior study of Hegg et al. (2018), were analyzed using the LASS and SS configurations. Prior data for each otolith was collected using a 30 μm spot size. Each otolith was ablated using parallel laser scans from the edge to the core of the otolith (Figure 1), perpendicular to the sulcus, in the same manner as prior studies on this population (Hegg et al. 2013, 2018, Chittaro et al. 2019). The limited size of the otolith cores necessitated testing at only three laser spot sizes. For LASS analysis, otoliths were ablated at laser spot sizes of 35, 25, and 15 μm. For SS analysis, otoliths were ablated at 25, 15, and 10 μm spot sizes, assuming increased sample at the detector in single stream mode. The new laser ablation system does not provide a 30 μm spot size, necessitating the use of 35 and 25 μm spots to bracket the prior data.

**Figure 1.**
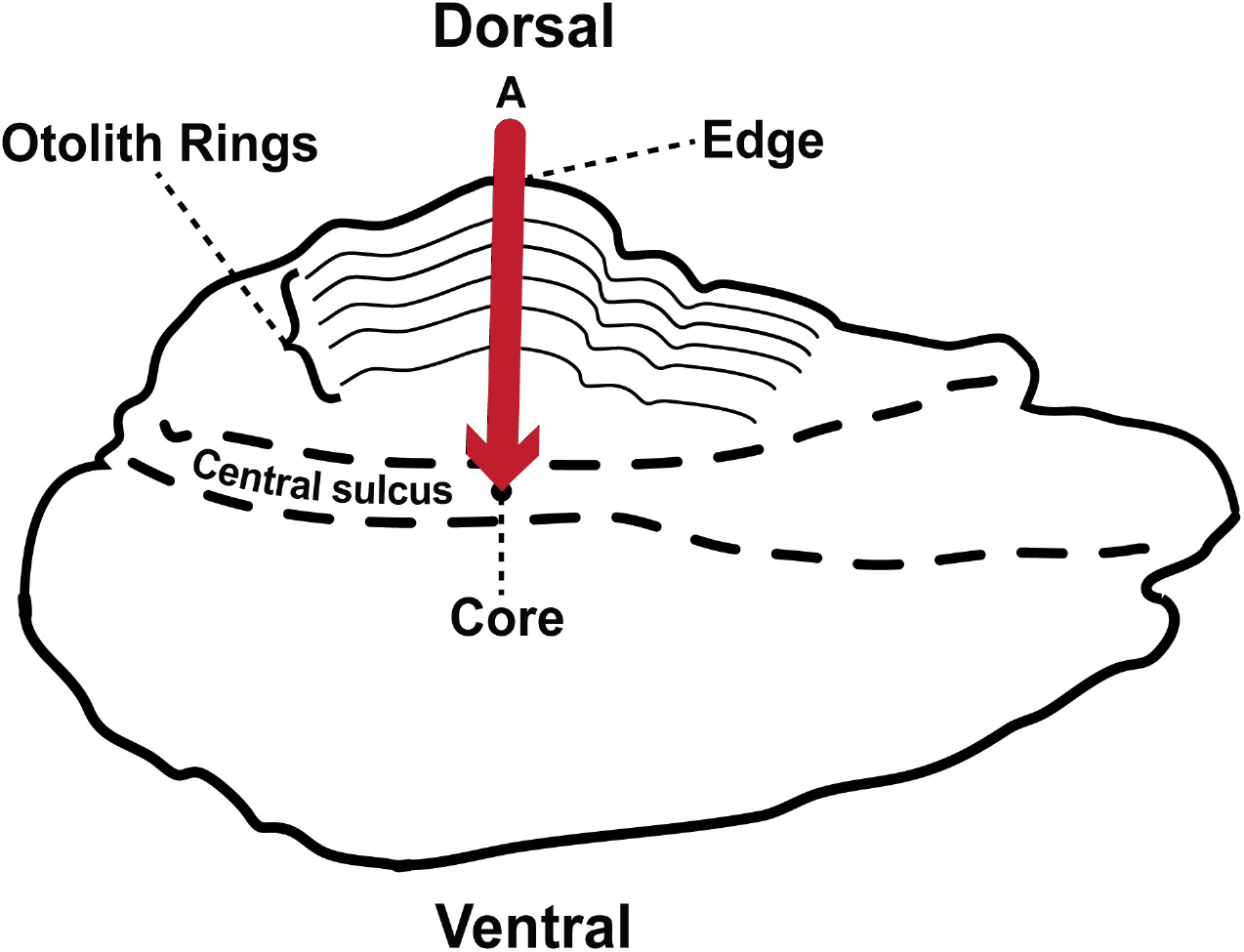
*Conceptual diagram of a Chinook salmon otolith, showing concentric ring structure. Laser ablation (A) proceeded from the otolith edge to the otolith core, perpendicular to the otolith sulcus. Data were then reversed to display chemical changes in chronological order from birth (Core) to death (Rim). Figure adapted from* Hegg *et al*. (2013)

### Instrument Settings

All work was performed in the Radiogenic Isotope and Geochronology Laboratory (RIGL) at Washington State University. Laser ablation analyses were done using both ‘traditional’ and LASS configurations. In the traditional configuration the laser ablation system (LA) is connected to a single inductively couple plasma mass spectrometer (LA-ICPMS). In LASS configuration, the ablated particles are ‘split’ such that the same sample volume is introduced to two independent mass spectrometers (Figure 2). In this study, a Teledyne 193nm ArF laser was coupled to a ThermoFisher Element2 sector-field ICPMS for determination of elemental concentrations, and a ThermoFisher Neptune Plus multi-collector ICPMS (MC-ICPMS) for determination of Sr isotope composition. Analyses were all done using a laser rastering speed of 10um/sec, with the laser operating at 20 Hz, and a fluence of ~4 J/cm^2^. A circular aperture was employed, and spot sizes ranged from 40 to 10 μm.

**Figure 2.**
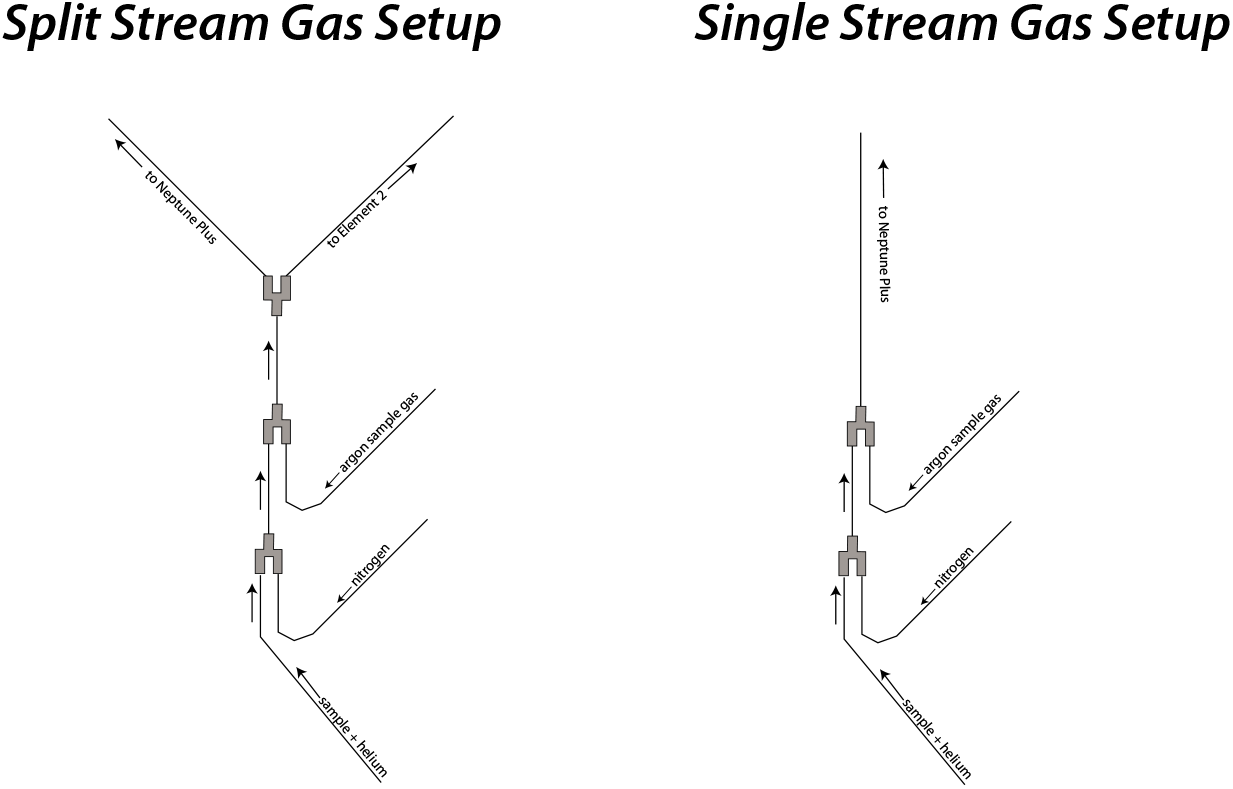
Conceptual diagram of gas and sample flows in the split stream (LASS) configuration (left) and single stream (right).

In both configurations the ablated sample aerosol is flushed from a two-volume laser cell using high-purity He for a carrier gas. Approximately 1m downstream from the laser cell, ~7-12 mL/min of nitrogen is introduced (dependent upon instrument configuration). Downstream of the nitrogen introduction, the “sample gas” from both the Element2 and NeptunePlus are added separately, before a final y-piece that splits the ablated particles along with the He-Ar-N carrier gas mixture to both mass spectrometers (Figure 2).

The instrument operating parameters are given in Table 1. The NeptunePlus MC-ICPMS is equipped with 9 Faraday detectors, which measure masses of Sr along with isobaric interference from Kr in the Ar supply, and Rb in the sample matrix (although this is almost always negligible in otoliths). In this study, we employ the combination of the ThermoFisher “Jet” sample cone, and “X” skimmer cone, which has been shown to offer the highest sensitivity, though the mass bias behavior is highly sensitive to both Ar and N gas flows (Fisher et al., 2020). We overcome this issue, yet take advantage of this high sensitivity cone configuration, by tuning gas flows prior to each session such that the mass-bias corrected ^87^Sr/^86^Sr is within 0.000150 of the accepted value for modern day ocean water 0.70918 (Kuznetsov et al. 2012, Mokadem et al. 2015, El Meknassi et al. 2018) on the basis of our in-house marine shell standard (0.709186, SD = 0.000077, n=535; Hegg, et al., 2018). Any residual offset from this canonical modern ocean water value is corrected by normalization to analyses of the MMS during the same analytical session. Both mass bias and Rb and Kr interferences corrections were performed for each 0.262 second integration. Mass bias was corrected using ^86^Sr/^88^Sr=0.1194. Interferences from ^87^Rb were monitored using ^85^Rb and corrected using ^87^Rb/^85^Rb = 0.38560. Interferences from ^84^Kr and ^88^Kr were monitored using ^83^Kr, and corrected using ^84^Kr/^83^Kr=4.95565 and ^86^Kr/^83^Kr =1.5026. The Sr isotope data are reduced using a customized ‘data reduction scheme’ (DRS; available from the second author) written for the Iolite software platform (Paton et al. 2011). For LASS data, we employ a bespoke version of this software specifically designed for viewing and data reduction of LASS analysis, described in detail by Fisher et al. (2017a).

**Table 1.**
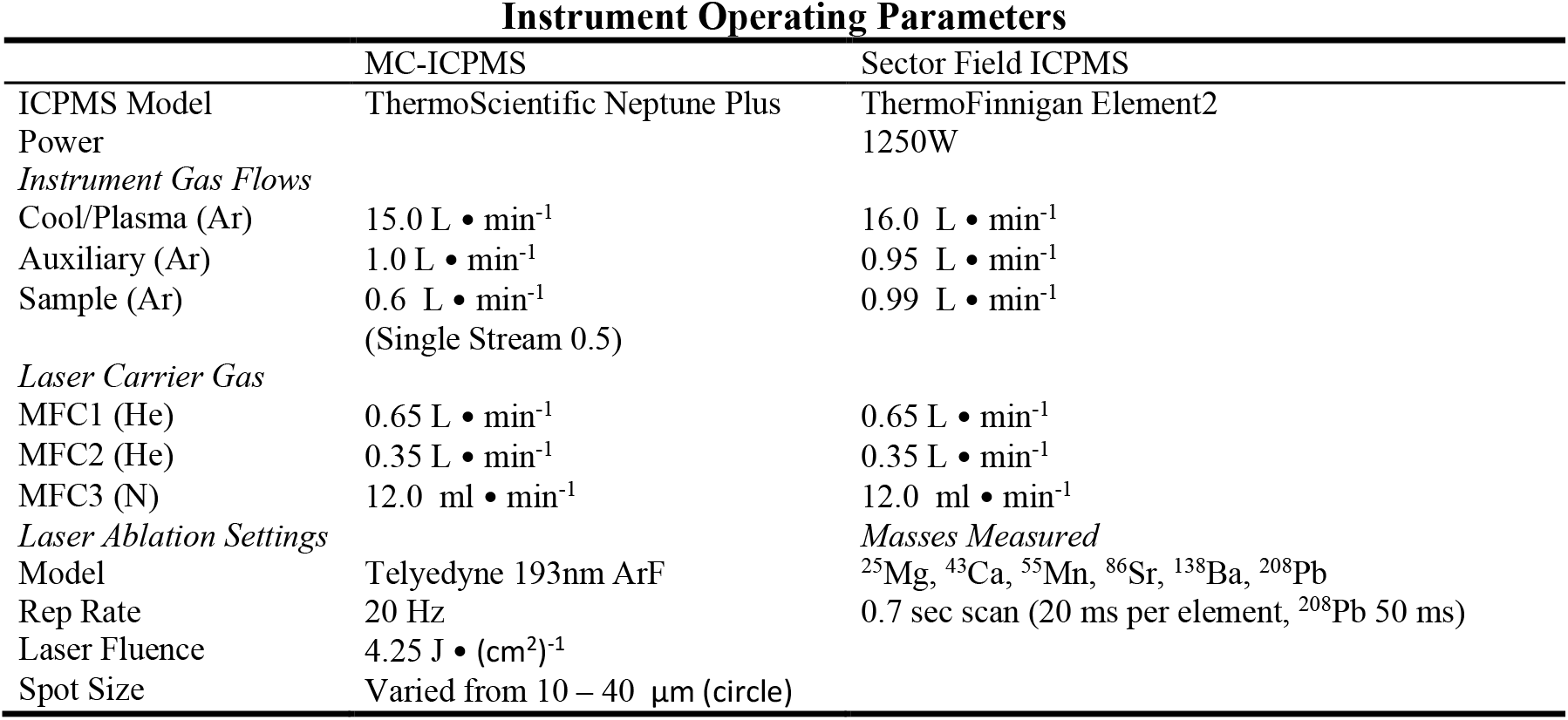
Instrument operating parameters for laser ablation analysis in the RIGL at WSU

The Element2 high resolution sector field ICPMS measured ^25^Mg, ^43^Ca, ^55^Mn ^138^Ba, and ^208^Pb. Each mass has 20 ms dwell time, with the exception of the 50 ms dwell time used for the low abundance ^208^Pb typically observed in otoliths and shells. While quadrupole ICPMS instruments offer very rapid scanning, their abundance sensitivity is notably lower than high-sensitivity sector-field instruments like the Element2, which allow robust determination of elements like Pb that occur in very low concentrations in otoliths and shells. Absolute concentrations are calibrated using the NIST 610 glass and also calculated using the Iolite software platform designed for LASS, with Ca as the internal standard. Otolith trace element contents are reported as elemental ratios to calcium, the internal standard. This is done to control for the variable ablation of sample, and because this ratio best represents the degree of incorporation of measured elements within the CaCO3 matrix of the otolith (Campana 1999).

### Statistical Analysis

Following data reduction to determine elemental concentrations and Sr isotope composition in the Iolite software package (Fisher et al. 2017a), otolith data were further reduced to determine the ^87^Sr/^86^Sr value during the rearing phase of each fish (Hegg et al. 2018). The fish analysed represent individuals from each of the four spawning areas within the study area, all rearing in the Snake River, and thus provide a realistic application of the method to fish with multiple life-histories within a large and heterogenous watershed. Further, the lower concentration of Sr in fresh water generally leads to higher uncertainty within this section of the otolith than in the period of marine growth during which ^87^Sr/^86^Sr is biologically uninformative.

To compare the effect of spot size and method (LASS versus Single Stream) on ^87^Sr/^86^Sr, a two-way ANOVA was performed on data from the modern marine shell standard with method and laser spot size as the grouping variables.

Reduction of prior data for the four comparison otoliths were done as described in Hegg et al. (2013, 2018) for ^87^Sr/^86^Sr. Trace element data were reduced using the elementR package for R (R Development Core Team 2011, Sirot et al. 2017) using published CRI values for the NIST 610 standard (Gagnon et al. 2008)

## Results

Standard errors increased with decreasing spot size for ^87^Sr/^86^Sr on the MMS standard (Figure 3A, Table 2). Spot sizes of 35 and 40 μm were comparable to the standard error of prior data at a 30 μm spot size on the same standard, but standard error increased.

**Table 2.**
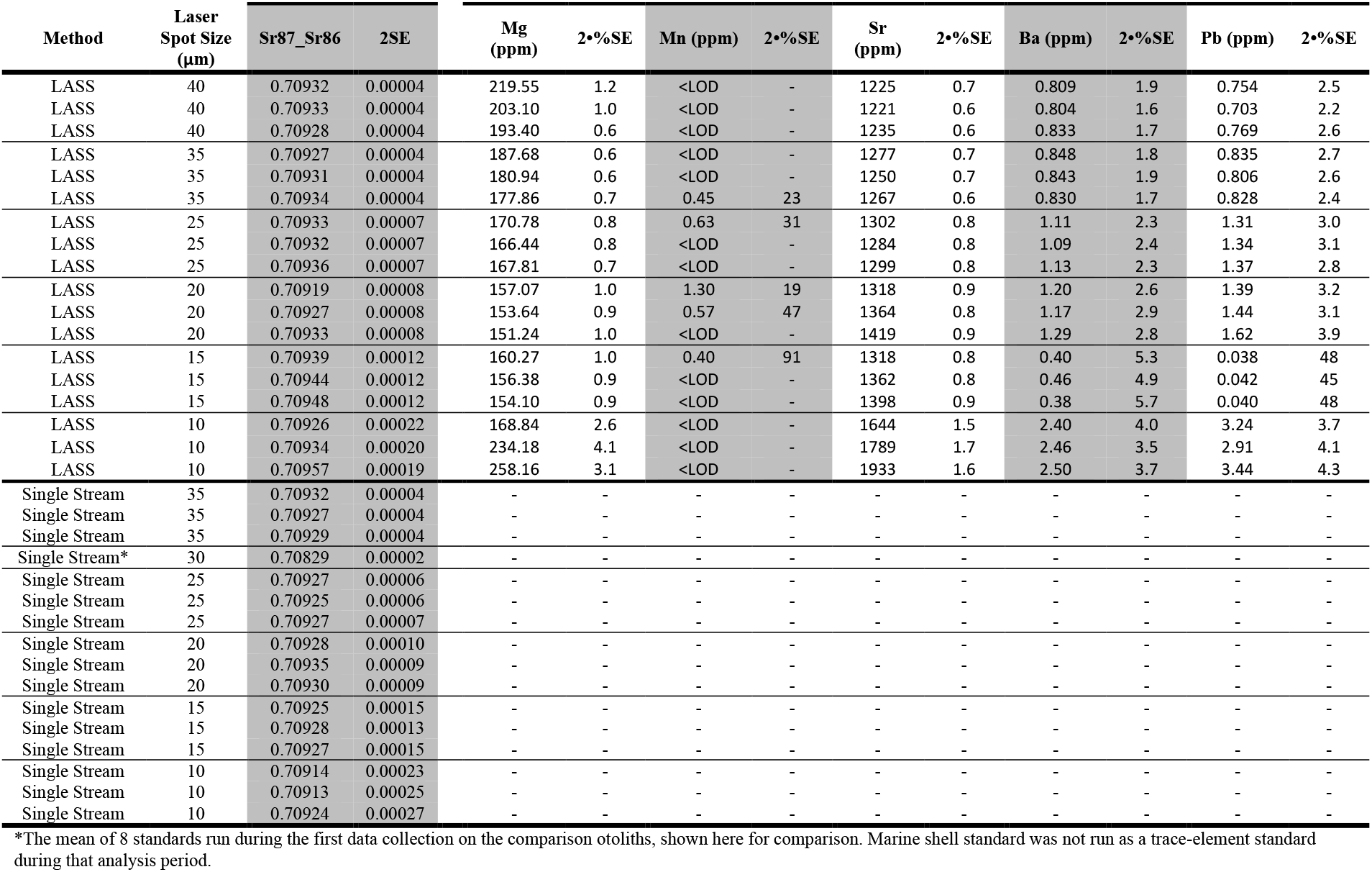
Analysis results of marine shell analytical standard at multiple spot sizes during LASS and SS.

**Figure 3.**
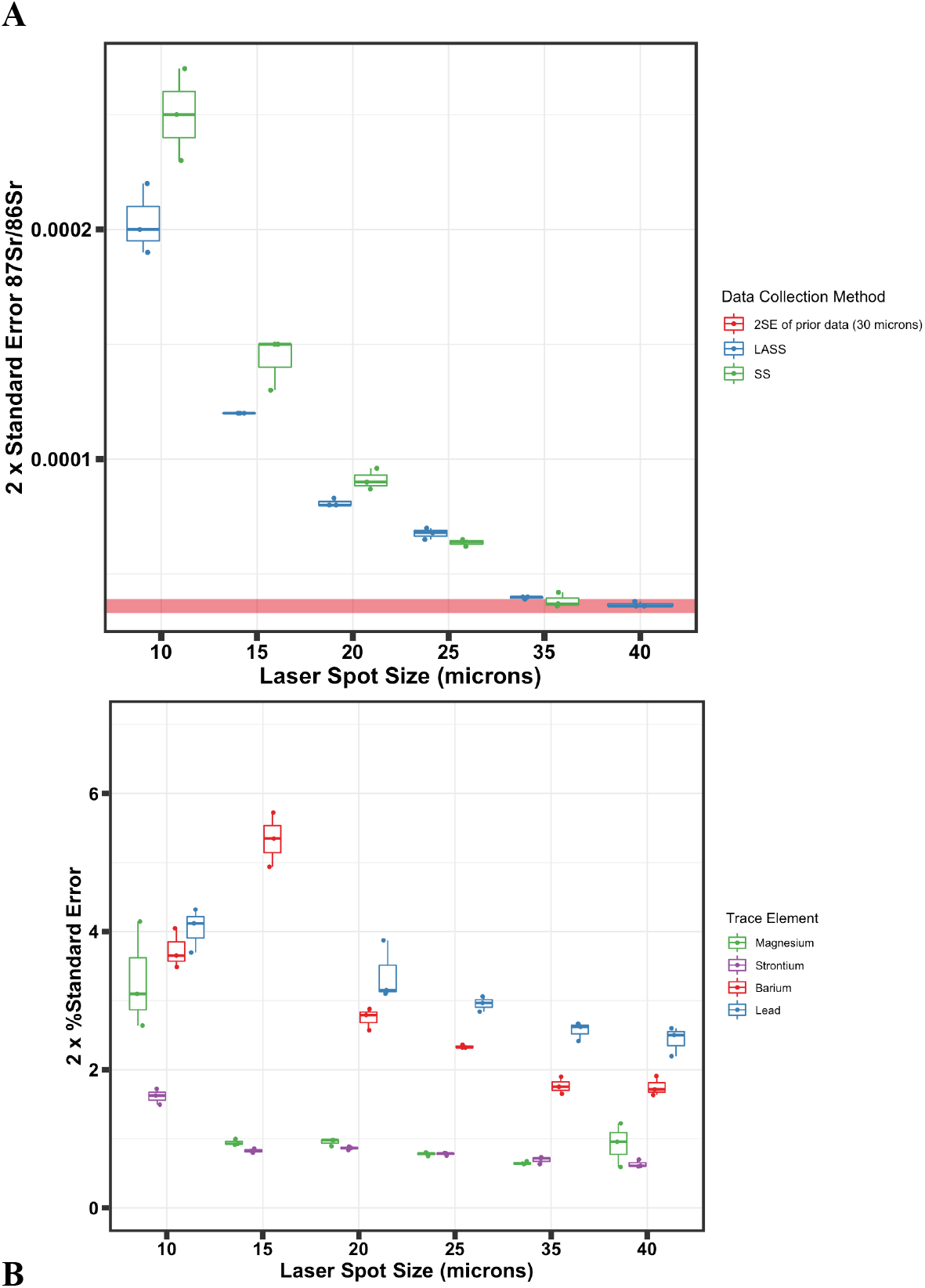
The standard error of strontium ratio measurements on a marine shell standard (**A**) decreases with increasing laser spot size in both LASS (blue) and single stream (green). The standard error of prior otolith work (red band) performed using a 30 μm spot is shown. The width of the band represents the mean and 2SE of 7 analyses of the same standard run during prior analysis of the 4 comparison otoliths. Percent standard error remained low for the most abundant isotopes, Sr and Mg down to 15 μm spot size. The least abundant isotopes, Ba and Pb, showed increases in uncertainty below 35 μm laser spot size (**B**).

Conversely, uncertainty in the simultaneous trace element analysis remained relatively stable, with uncertainty for only the least abundant elements increasing marginally down to 15 μm spot size after which standard errors markedly increased in the least abundant elements of lead and barium, and finally large increases in percent standard error for strontium and magnesium at 10 μm laser spot size (Figure 3B, Table 2).

Lead and manganese were present in the marine shell at very low levels. Manganese values were largely below detection limit and were therefore removed from further data analysis. Lead values were extremely low but detectable.

Mean strontium isotope ratios of the MMS, corrected for mass bias and Kr interference were compared across all spot sizes using a two-way ANOVA to compare the effect of the method (LASS vs. Single Stream), laser spot size, and their interaction term. These results were uncorrected to the canonical ocean signature since this would require a correction using the MMS itself. Method was a significant model variable (p = 0.0014), laser spot size was not significant (p = 0.3509), indicating a significant difference in mean ^87^Sr/^86^Sr between LASS and single stream but not between spot sizes. The interaction between measurement method with spot size was significant (p = 0.0099), indicating differing mean ^87^Sr/^86^Sr was related to an interaction between method and laser spot size. This interaction is evident in Figure 4, with ^87^Sr/^86^Sr values remaining similar between the two methods until laser spot size falls to 15 μm, at which point the methods diverge, with single stream resulting in lower values and LASS resulting in higher values.

**Figure 4.**
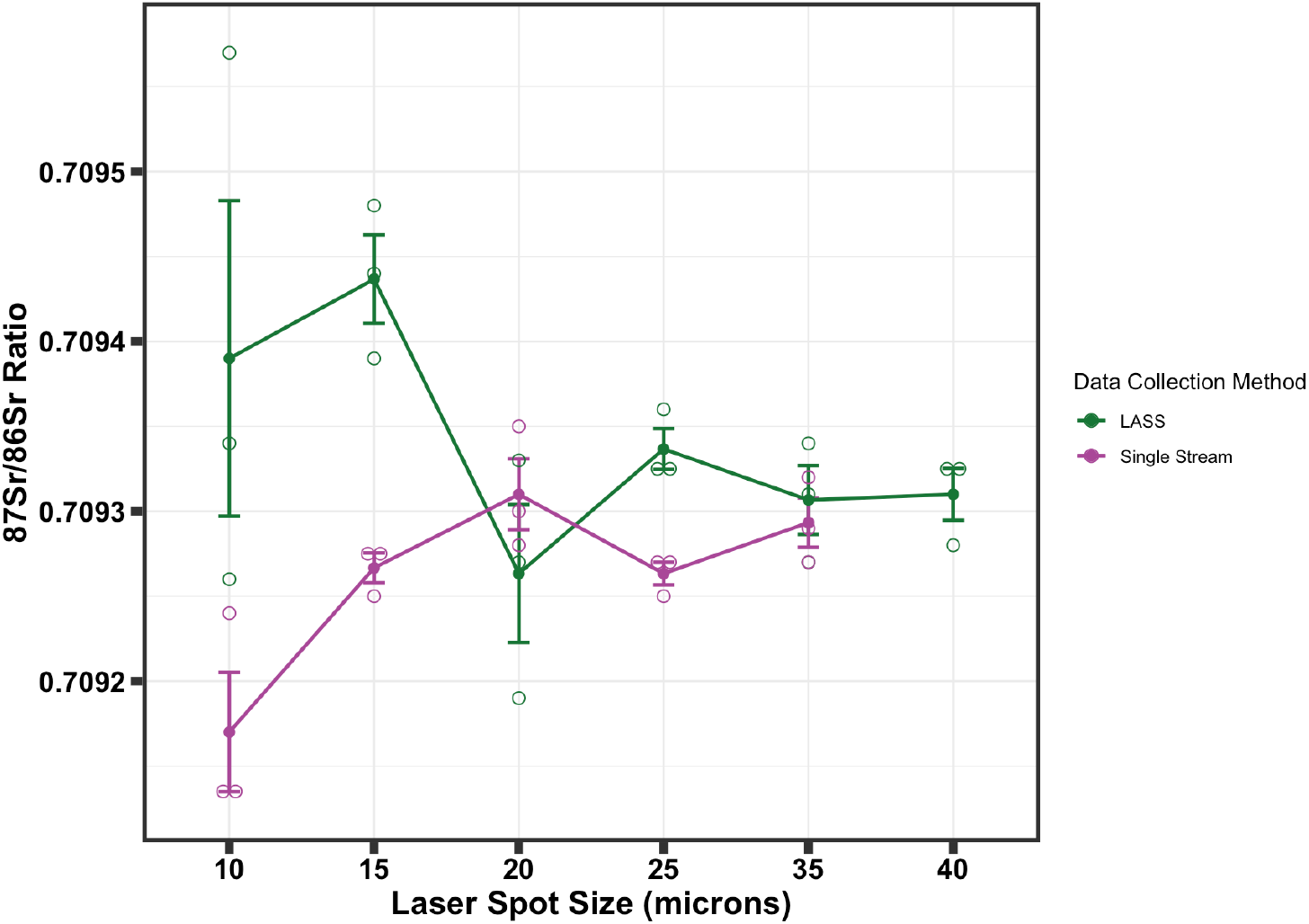
Comparison of ^87^Sr/^86^Sr precision on the marine shell standard at varying spot sizes between LASS and single stream data collection.

Analysis of fish otoliths shows the effect of spot size on ^87^Sr/^86^Sr uncertainty. After correcting the SS data to match the lower integration rate of the LASS data reduction in Iolite, the uncertainties across each otolith were similar in LASS and single stream configurations. Figure 5 shows Sr isotope data from a representative otolith. The juvenile section of the salmon otoliths (between 0 and ~1100 μm from the otolith core in Figure 5) contain lower strontium concentrations due to their formation in fresh water, resulting in higher uncertainties across these sections of the otolith. For this reason, 25 μm spot sizes began to show data with precision outside the optimal range for otolith analysis (Figure 5), with smaller spot sizes showing increasingly higher uncertainty.

**Figure 5.**
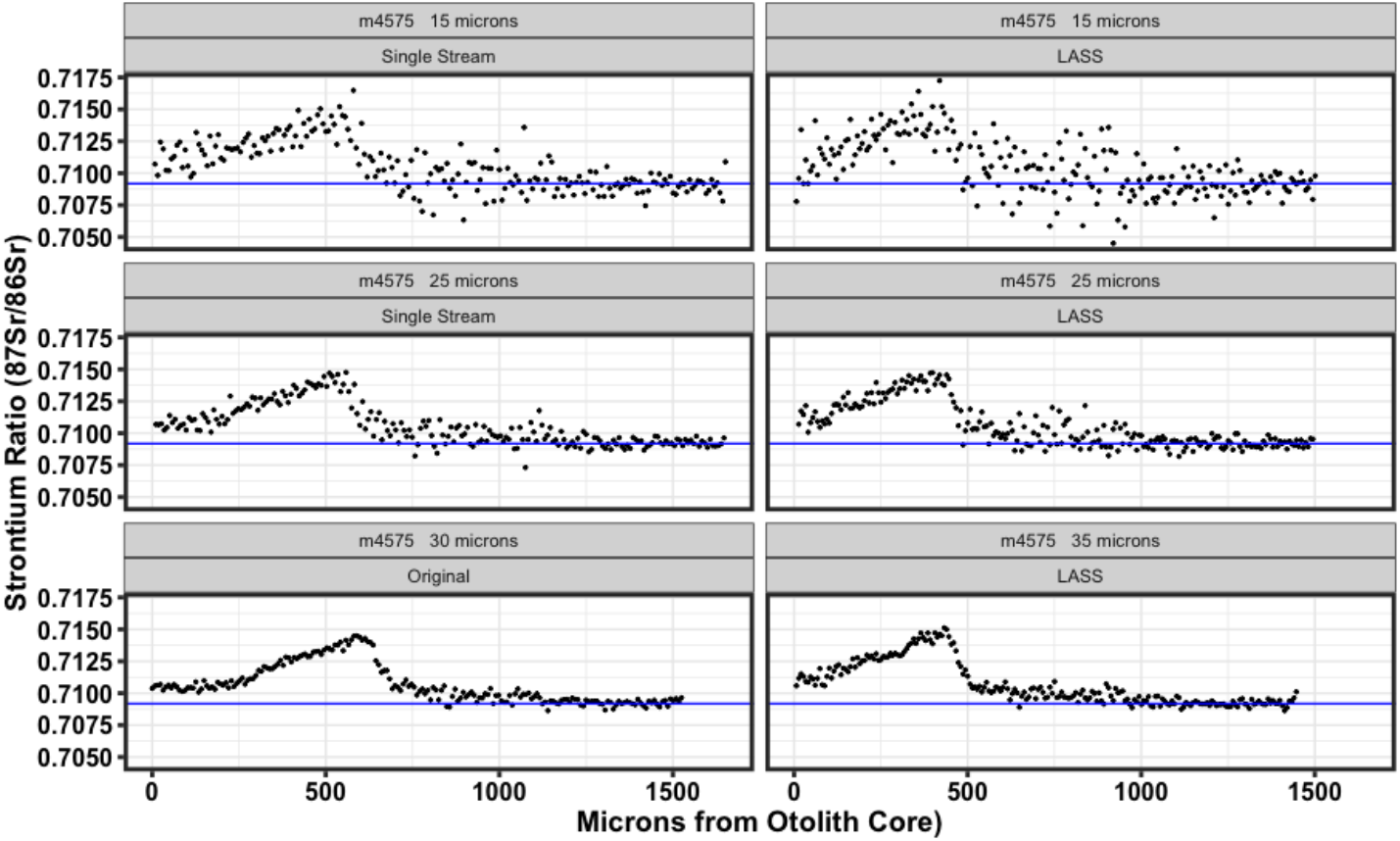
Comparison of strontium ratio data analysed at multiple laser spot sizes (rows) and using single-stream (column 1) and LASS (column 2) on a continuous transect across the same otolith using a Thermo Neptune Plus and Telyedyne 193nm ArF laser combination. Prior data collected on a Thermo Neptune and New Wave UP-213 laser with a 30 μm spot size are shown (row 3, column 1). Single Stream data have been downsampled by averaging every 3 integrations to better match LASS data which were output at a lower integration rate.

The rearing section of each otolith, identified through prior research by Hegg et al. (2018), was averaged from both the LASS and single stream otolith data to provide a real world comparison of performance of the methods (Table 3). These periods are of differing lengths depending on the life history of the individual fish. Further, the parallel position of each subsequent laser track on the otolith causes contraction and dilation of the pattern across the otolith, resulting in slightly different samples averaged from each fish.

**Table 3.**
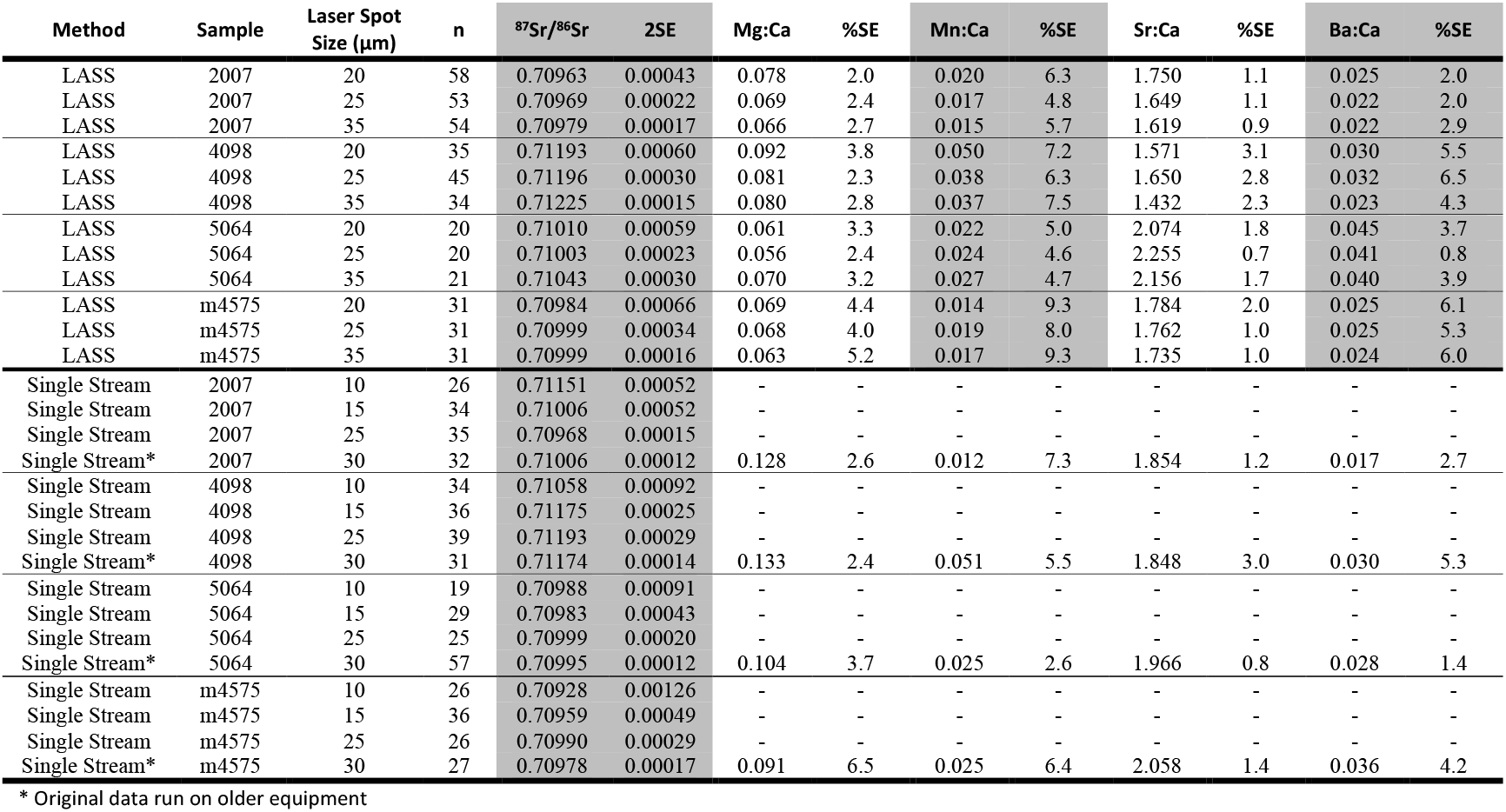
Analysis of the rearing period of four adult Fall Chinook salmon otoliths using LASS and Single Stream methods. Trace element ratios are reported in ppm x 1000.

| Method | Sample | Laser Spot Size (µm) | n | <sup>87</sup>Sr/<sup>86</sup>Sr | 2SE | Mg:Ca | %SE | Mn:Ca | %SE | Sr:Ca | %SE | Ba:Ca | %SE |
| --- | --- | --- | --- | --- | --- | --- | --- | --- | --- | --- | --- | --- | --- |
| LASS | 2007 | 20 | 58 | 0.70963 | 0.00043 | 0.078 | 2.0 | 0.020 | 6.3 | 1.750 | 1.1 | 0.025 | 2.0 |
| LASS | 2007 | 25 | 53 | 0.70969 | 0.00022 | 0.069 | 2.4 | 0.017 | 4.8 | 1.649 | 1.1 | 0.022 | 2.0 |
| LASS | 2007 | 35 | 54 | 0.70979 | 0.00017 | 0.066 | 2.7 | 0.015 | 5.7 | 1.619 | 0.9 | 0.022 | 2.9 |
| LASS | 4098 | 20 | 35 | 0.71193 | 0.00060 | 0.092 | 3.8 | 0.050 | 7.2 | 1.571 | 3.1 | 0.030 | 5.5 |
| LASS | 4098 | 25 | 45 | 0.71196 | 0.00030 | 0.081 | 2.3 | 0.038 | 6.3 | 1.650 | 2.8 | 0.032 | 6.5 |
| LASS | 4098 | 35 | 34 | 0.71225 | 0.00015 | 0.080 | 2.8 | 0.037 | 7.5 | 1.432 | 2.3 | 0.023 | 4.3 |
| LASS | 5064 | 20 | 20 | 0.71010 | 0.00059 | 0.061 | 3.3 | 0.022 | 5.0 | 2.074 | 1.8 | 0.045 | 3.7 |
| LASS | 5064 | 25 | 20 | 0.71003 | 0.00023 | 0.056 | 2.4 | 0.024 | 4.6 | 2.255 | 0.7 | 0.041 | 0.8 |
| LASS | 5064 | 35 | 21 | 0.71043 | 0.00030 | 0.070 | 3.2 | 0.027 | 4.7 | 2.156 | 1.7 | 0.040 | 3.9 |
| LASS | m4575 | 20 | 31 | 0.70984 | 0.00066 | 0.069 | 4.4 | 0.014 | 9.3 | 1.784 | 2.0 | 0.025 | 6.1 |
| LASS | m4575 | 25 | 31 | 0.70999 | 0.00034 | 0.068 | 4.0 | 0.019 | 8.0 | 1.762 | 1.0 | 0.025 | 5.3 |
| LASS | m4575 | 35 | 31 | 0.70999 | 0.00016 | 0.063 | 5.2 | 0.017 | 9.3 | 1.735 | 1.0 | 0.024 | 6.0 |
| Single Stream | 2007 | 10 | 26 | 0.71151 | 0.00052 | - | - | - | - | - | - | - | - |
| Single Stream | 2007 | 15 | 34 | 0.71006 | 0.00052 | - | - | - | - | - | - | - | - |
| Single Stream | 2007 | 25 | 35 | 0.70968 | 0.00015 | - | - | - | - | - | - | - | - |
| Single Stream* | 2007 | 30 | 32 | 0.71006 | 0.00012 | 0.128 | 2.6 | 0.012 | 7.3 | 1.854 | 1.2 | 0.017 | 2.7 |
| Single Stream | 4098 | 10 | 34 | 0.71058 | 0.00092 | - | - | - | - | - | - | - | - |
| Single Stream | 4098 | 15 | 36 | 0.71175 | 0.00025 | - | - | - | - | - | - | - | - |
| Single Stream | 4098 | 25 | 39 | 0.71193 | 0.00029 | - | - | - | - | - | - | - | - |
| Single Stream* | 4098 | 30 | 31 | 0.71174 | 0.00014 | 0.133 | 2.4 | 0.051 | 5.5 | 1.848 | 3.0 | 0.030 | 5.3 |
| Single Stream | 5064 | 10 | 19 | 0.70988 | 0.00091 | - | - | - | - | - | - | - | - |
| Single Stream | 5064 | 15 | 29 | 0.70983 | 0.00043 | - | - | - | - | - | - | - | - |
| Single Stream | 5064 | 25 | 25 | 0.70999 | 0.00020 | - | - | - | - | - | - | - | - |
| Single Stream* | 5064 | 30 | 57 | 0.70995 | 0.00012 | 0.104 | 3.7 | 0.025 | 2.6 | 1.966 | 0.8 | 0.028 | 1.4 |
| Single Stream | m4575 | 10 | 26 | 0.70928 | 0.00126 | - | - | - | - | - | - | - | - |
| Single Stream | m4575 | 15 | 36 | 0.70959 | 0.00049 | - | - | - | - | - | - | - | - |
| Single Stream | m4575 | 25 | 26 | 0.70990 | 0.00029 | - | - | - | - | - | - | - | - |
| Single Stream* | m4575 | 30 | 27 | 0.70978 | 0.00017 | 0.091 | 6.5 | 0.025 | 6.4 | 2.058 | 1.4 | 0.036 | 4.2 |

\* Original data run on older equipment

Strontium isotope ratio measurements within the rearing section of each otolith exhibited an order of magnitude more uncertainty than the marine shell due to the decreased strontium concentration of fresh water and the smaller number of integrations used to calculate the average. Despite this, strontium isotope ratios across each laser spot size and between LASS and single stream methods were in agreement within each otolith (Figure 6). Mn/Ca and Ba/Ca ratios were consistently slightly lower than data from the original study. The reason for this is unknown but the low standard errors and consistent results during LASS indicate that this is unlikely to be related to the LASS configuration. The limited size of the otolith cores made it impossible to make multiple analyses at each laser spot size for statistical comparison.

**Figure 6.**
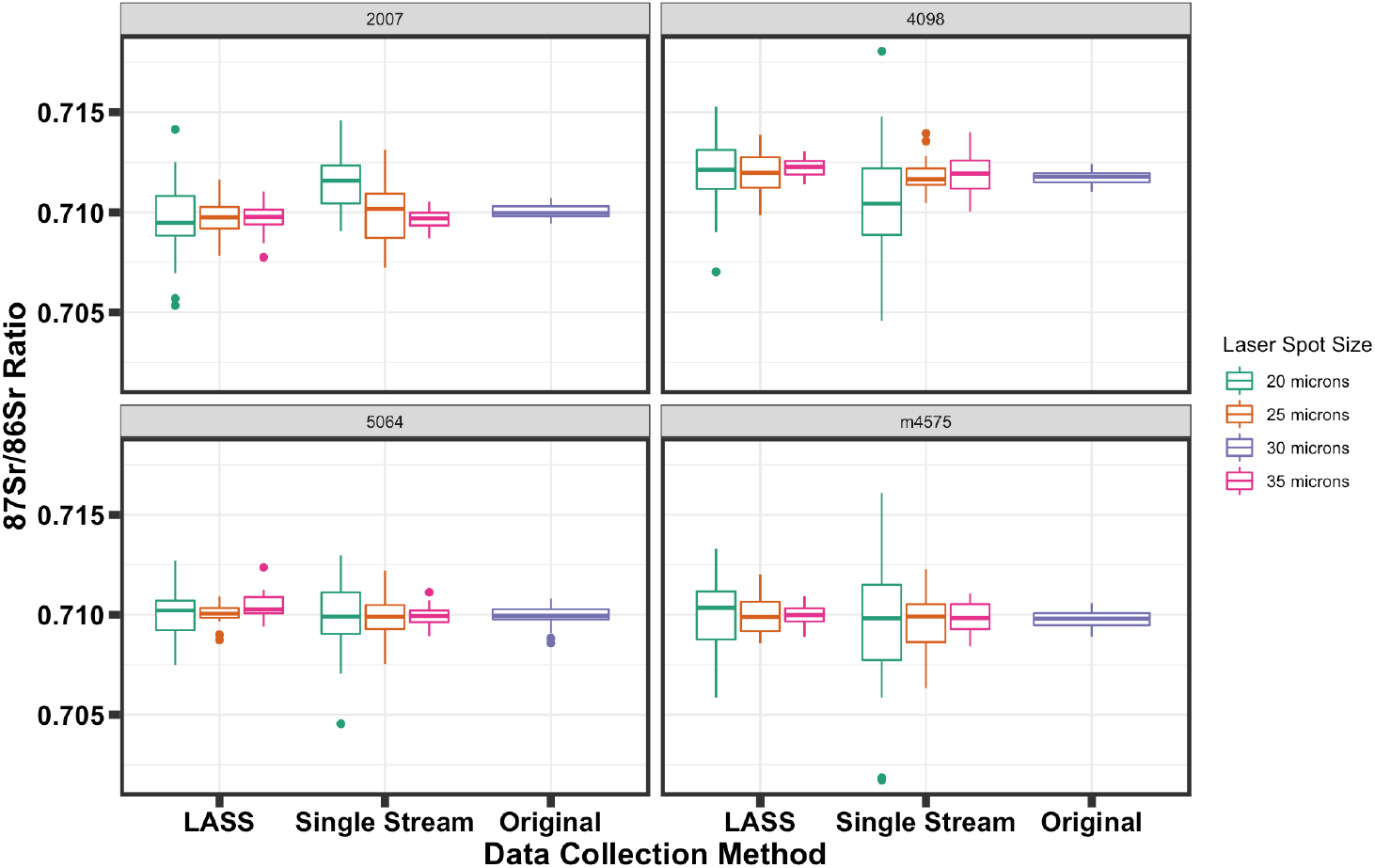
Strontium isotope ratios measured in the rearing sections of four salmon otoliths using LASS, and single stream collection methods and at differing laser spot sizes. Original data from a prior study are shown for comparison.

## Discussion

The increase in analytical precision and throughput of LA-ICPMS methodology has spurred diverse new methods by which chronological hard structures can be used to reconstruct past events. Using corals as recorders of ocean chemistry (Rüggeberg et al. 2008, Mallela et al. 2011), stalactites to track monsoon rains and glaciation (Zhou et al. 2009, Griffiths et al. 2010), and otoliths to understand migration of fish (Hamann and Kennedy 2012, Seeley et al. 2015) multiple techniques have emerged which can be used simultaneously to reconstruct multiple histories from a single sample.

Walther (2019) highlights the importance of multiple data streams from otoliths to interpret the complexity of fish life history in otoliths. This is also true of similar chronological structures from bivalve shells, mammal teeth, or stalactites. Often, many processes are recorded simultaneously by different tracers and the totality gives an improved understanding relative to any single tracer. As technology and throughput have improved in the study of chronological hard structures, the desire to utilize multiple analytes within the same sample has increased as well. These multiple tracers provide multiple simultaneous perspectives on climate, migration, growth, hydrology, or temperature that greatly increase our ability to reconstruct and interpret data.

However, because these structures are chronological, each element or isotopic ratio must be well aligned with the others to provide meaning through time. This can be difficult when combining isotopic and trace element data from multiple instruments with slightly variable laser track lengths and different integration times, potentially introducing additional uncertainty. Using the LASS data reduction methods developed for Iolite (Fisher et al. 2017b) with simultaneous collection of data from the same laser track eliminates these alignment errors. This results in output data that are matched to the same integration time and thus easier to analyze as well as potentially doubling throughput.

Because the sample is split between two ICPMS instruments during LASS analysis, a spot size large enough to provide sufficient sample to each detector is required to maintain precision. Validating LA-MC-ICPMS for ^87^Sr/^86^Sr analysis Barnett-Johnson (2005) reported external error of ±0.00003 (2SE) with an 80 μm spot size on marine carbonate. Our analysis on the marine shell standard indicates that similar levels of precision (±0.00004) can be achieved with 35 and 40 μm spot sizes on current equipment, with 25 μm spot sizes yielding only slightly higher uncertainties (Table 2). Precision was less dependent on spot size for trace-element analysis during LASS, where standard error showed a moderate increase down to 20 μm laser spot diameter before becoming highly imprecise.

Laser spot size was not significant in the ANOVA analysis, however visual inspection of the results indicates that accuracy deteriorated at the smallest spot sizes (Figure 4). The best results were obtained at 35 and 40 μm spot sizes; at spot sizes smaller than 20 μm accuracy degraded. The reason for the difference in ^87^Sr/^86^Sr results between LASS and single stream methods at 25 μm is unknown but could be related to variation across the marine shell given that precision of these measurements was high, or due to the effects of using the new high-sensitivity cone combination in the presence of nitrogen. However, this does highlight the significant effect of method in the ANOVA of the MMS standard. In this study LASS tended to err with a positive bias while single stream erred low, resulting in the significant ANOVA result.

Data from the salmon otoliths indicate that the LASS methodology performed similarly to single stream for ^87^Sr/^86^Sr analysis, and accuracy compared well to prior results (Figure 6). In the Snake River, where these fish were captured, tributary ^87^Sr/^86^Sr varies at the scale of 1×10^−3^ (Hegg et al. 2013). Given this, data for the example otolith in Figure 4 is still useful for classifying fish location down to a 25 μm laser spot size, even with the small sample size following clipping the rearing section of the transect. However, for systems with lower variability in background signature larger spot sizes or longer scans along the otolith rings might be needed. Regardless, these findings demonstrate that LASS analyses can produce results with precision akin to traditional dedicated (single stream) methods.

Trace element results for the rearing sections of the four otoliths are difficult to interpret beyond the broadest scale. While ^87^Sr/^86^Sr is stable during the rearing period other elements do not necessarily remain stable and their signatures in all four fish exhibited changes during the rearing period. Thus, the average of any given trace element is heavily affected by variations in the clipped portion. Because each laser track was performed parallel to the others, the curved nature of the otolith rings resulted in minor expansion and compression of the chemical patterns between runs, which are visible in the example otolith in Figure 4. This resulted in unavoidable differences in the clipped portion and the inability to easily compare their values in anything but the broadest sense. Because trace elements are affected both by environment and ontogeny, during the course of an otolith study the magnitude of the elemental ratios would cluster for fish following similar life histories, however in this context the exact analytical accuracy and precision is difficult to measure.

The marine shell standard used is a natural shell and is not intended as a suitable monitor for trace element accuracy or precision. Variation in trace elements exists across the surface of the standard, as has been demonstrated in many marine bivalves (Butler and Schöne 2017, Markulin et al. 2019). A homogenous carbonate standard was not available during this study. Therefore, we recommend that future work to validate this method, with respect to trace elements, should examine the stability of trace element ratios using one of the available homogenous carbonate standards (Weber et al. 2018, Jochum et al. 2019). The sensitivity of the Element2 sector field ICP-MS, and the low standard error of abundant trace elements for LASS runs across the majority of laser spot sizes, indicate that ^87^Sr/^86^Sr, not trace-elements, is the method most prone to error at small spot sizes within the LASS system.

The ability to generate and easily reduce simultaneously determined Sr isotope ratio and trace element contents of samples represents a significant advance in the collection of chronological transect data from carbonate samples. The need for consistent reduction methods has been noted by multiple authors (Sirot et al. 2017, Willmes et al. 2018) and the addition of LASS methods in this study presents an additional approach to reduce alignment and data reduction errors. The method detailed here shows that accuracy and precision of ^87^Sr/^86^Sr ratios can be achieved at smaller spot sizes than in prior attempts at LASS on fish otoliths (Prohaska et al. 2016). Further, these results can be achieved while outputting time and integration matched data using novel Iolite reduction software, significantly shortening the reduction and analysis time for multiple data streams. While the elements and isotope ratios demonstrated here are limited, the data reduction scheme is scalable to multiple trace-element ratios and isotope systems, as well as the use of multiple ICPMS platforms (Fisher et al. 2017a).

## Acknowledgements

Thank you to B. Kennedy and the University of Idaho LIFE lab for manuscript edits and use of the otolith samples and prior data. Thank you to B. Hamann, E. Benson, and J. Caisman for help in preparation and analysis of the original otolith samples.

## References Cited

Andrus, C. F. T. 2011. Shell midden sclerochronology. Quaternary Science Reviews 30:2892–2905.

Barnett-Johnson, R., F. C. Ramos, C. B. Grimes, and R. B. MacFarlane. 2005. Validation of Sr isotopes in otoliths by laser ablation multicollector inductively coupled plasma mass spectrometry (LA-MC-ICPMS): opening avenues in fisheries science applications. Canadian Journal of Fisheries and Aquatic Sciences 62:2425–2430.

Bath, G. E., S. R. Thorrold, C. M. Jones, S. E. Campana, J. W. McLaren, and J. W. H. Lam. 2000. Strontium and barium uptake in aragonitic otoliths of marine fish. Geochimica et Cosmochimica Acta 64:1705–1714.

Begg, G. A., S. E. Campana, A. J. Fowler, and I. M. Suthers. 2005. Otolith research and application: Current directions in innovation and implementation. Pages 477–483 Marine and Freshwater Research.

Brennan, S. R., and D. E. Schindler. 2017. Linking otolith microchemistry and dendritic isoscapes to map heterogeneous production of fish across river basins: Ecological Applications 27:363–377.

Butler, P. G., and B. R. Schöne. 2017, January 1. New research in the methods and applications of sclerochronology. Elsevier B.V.

Campana, S. 1999. Chemistry and composition of fish otoliths:pathways, mechanisms and applications. Marine Ecology Progress Series 188:263–297.

Campana, S. E., and S. R. Thorrold. 2001. Otoliths, increments, and elements: keys to a comprehensive understanding of fish populations? Canadian Journal of Fisheries and Aquatic Sciences 58:30–38.

Campana, S. E., S. R. Thorrold, C. M. Jones, D. Günther, M. Tubrett, H. Longerich, S. Jackson, N. M. Halden, J. M. Kalish, P. Piccoli, H. de Pontual, H. Troadec, J. Panfili, D. H. Secor, K. P. Severin, S. H. Sie, R. Thresher, W. Teesdale, J. L. Campbell, S. Campana, S. Thorrold, C. Jones, M. Tubrett, H. Longerich, S. Jackson, N. Halden, J. Kalish, H. de Pontual, H. Troadec, J. Panfili LASAA, D. Secor, K. Severin, S. Sie Heavy, R. Thresher, and J. Campbell. 1997. Comparison of accuracy, precision, and sensitivity in elemental assays of fish otoliths using the electron microprobe, proton-induced X-ray emission, and laser ablation inductively coupled plasma mass spectrometry. Page Aquat. Sci.

Chittaro, P. M., J. C. Hegg, B. P. Kennedy, L. A. Weitkamp, L. L. Johnson, C. Bucher, and R. W. Zabel. 2019. Juvenile river residence and performance of Snake River fall Chinook salmon. Ecology of Freshwater Fish 28:396–410.

Crook, D. A., J. I. Macdonald, D. G. McNeil, D. M. Gilligan, M. Asmus, R. Maas, J. Woodhead, and B. Gillanders. 2013. Recruitment sources and dispersal of an invasive fish in a large river system as revealed by otolith chemistry analysis. Canadian Journal of Fisheries and Aquatic Sciences 70:953–963.

Fairchild, I. J., A. Baker, A. Asrat, D. Domínguez-Villar, J. Gunn, A. Hartland, and D. Lowe. 2012. Speleothem Science: From Process to Past Environments. Page Speleothem Science: From Process to Past Environments.

Fisher, C. M., C. Paton, D. G. Pearson, C. Sarkar, Y. Luo, D. B. Tersmette, and T. Chacko. 2017a. Data Reduction of Laser Ablation Split-Stream (LASS) Analyses Using Newly Developed Features Within Iolite: With Applications to Lu-Hf + U-Pb in Detrital Zircon and Sm-Nd +U-Pb in Igneous Monazite. Geochemistry, Geophysics, Geosystems 18:4604–4622.

Fisher, C. M., C. Paton, D. G. Pearson, C. Sarkar, Y. Luo, D. B. Tersmette, and T. Chacko. 2017b. Data Reduction of Laser Ablation Split-Stream (LASS) Analyses Using Newly Developed Features Within Iolite: With Applications to Lu-Hf + U-Pb in Detrital Zircon and Sm-Nd +U-Pb in Igneous Monazite. Geochemistry, Geophysics, Geosystems 18:4604–4622.

Fisher, C. M., J. D. Vervoort, and S. A. DuFrane. 2014. Accurate Hf isotope determinations of complex zircons using the “laser ablation split stream” method. Geochemistry, Geophysics, Geosystems 15:121-139.

Füllenbach, C. S., B. R. Schöne, and R. Mertz-Kraus. 2015. Strontium/lithium ratio in aragonitic shells of Cerastoderma edule (Bivalvia) - A new potential temperature proxy for brackish environments. Chemical Geology 417:341–355.

Gagnon, J. E., B. J. Fryer, I. M. Samson, and A. E. Williams-Jones. 2008. Quantitative analysis of silicate certified reference materials by LA-ICPMS with and without an internal standard.

Garcez, R. C. S., R. Humston, D. Harbor, and C. E. C. Freitas. 2014. Otolith geochemistry in young-of-the-year peacock bass Cichla temensis for investigating natal dispersal in the Rio Negro (Amazon - Brazil) river system. Ecology of Freshwater Fish:n/a-n/a.

Goudie, D. J., C. M. Fisher, J. M. Hanchar, J. L. Crowley, and J. C. Ayers. 2014. Simultaneous in situ determination of U-Pb and Sm-Nd isotopes in monazite by laser ablation ICP-MS. Geochemistry, Geophysics, Geosystems 15:2575–2600.

Griffiths, M. L., R. N. Drysdale, M. K. Gagan, S. Frisia, J. xin Zhao, L. K. Ayliffe, W. S. Hantoro, J. C. Hellstrom, M. J. Fischer, Y. X. Feng, and B. W. Suwargadi. 2010. Evidence for Holocene changes in Australian-Indonesian monsoon rainfall from stalagmite trace element and stable isotope ratios. Earth and Planetary Science Letters 292:27–38.

Hamann, E. J., and B. P. Kennedy. 2012. Juvenile dispersal affects straying behaviors of adults in a migratory population. Ecology 93:733–740.

Hegg, J. C., T. Giarrizzo, and B. P. Kennedy. 2015. Diverse Early Life-History Strategies in Migratory Amazonian Catfish: Implications for Conservation and Management. Plos One 10.

Hegg, J. C., B. Graves, and C. M. Fisher. 2019. Sawfish, Read in Tooth and Saw: rostral teeth as endogenous chemical records of movement and life-history in a critically endangered species. bioRxiv:753293.

Hegg, J. C., B. P. Kennedy, and P. M. Chittaro. 2018. What did you say about my mother? The complexities of maternally derived chemical signatures in otoliths. Canadian Journal of Fisheries and Aquatic Sciences 14:1–14.

Hegg, J. C., B. P. Kennedy, P. M. Chittaro, and R. W. Zabel. 2013. Spatial structuring of an evolving life-history strategy under altered environmental conditions. Oecologia 172:1017–1029.

Hendy, C. H., and A. T. Wilson. 1968. Palaeoclimatic data from speleothems. Nature 219:48–51.

Hobson, K. A. 1999. Tracing origins and migration of wildlife using stable isotopes: a review. Oecologia 120:314–326.

Hobson, K. A., R. Barnett-Johnson, and T. Cerling. 2010. Using isoscapes to track animal migration. Pages 273–298 Isoscapes: understanding movement, pattern, and process on Earth through isotope mapping. Springer Verlag.

Hudson, J. H., E. A. Shinn, R. B. Halley, and B. Lidz. 1976. Sclerochronology: A tool for interpreting past environments. Geology 4:361–364.

Jacques, P., D. P. Helene, T. Herve, and W. P. J. 2002. Manual of fish Sclerochronology. Ifremer.

Jochum, K. P., D. Garbe-Schönberg, M. Veter, B. Stoll, U. Weis, M. Weber, F. Lugli, A. Jentzen, R. Schiebel, J. A. Wassenburg, D. E. Jacob, and G. H. Haug. 2019. Nano-Powdered Calcium Carbonate Reference Materials: Significant Progress for Microanalysis? Geostandards and Geoanalytical Research 43:595–609.

Kennedy, B. P., C. L. Folt, J. D. Blum, and C. P. Chamberlain. 1997. Natural isotope markers in salmon. Nature 387:766–767.

Kennedy, B. P., A. Klaue, J. D. Blum, C. L. Folt, and K. H. Nislow. 2002. Reconstructing the lives of fish using Sr isotopes in otoliths. Canadian Journal of Fisheries and Aquatic Sciences 59:925–929.

Kohn, M. J., J. Morris, and P. Olin. 2013. Trace element concentrations in teeth–a modern Idaho baseline with implications for archeometry, forensics, and palaeontology. Journal of Archaeological Science 40:1689–1699.

Kuznetsov, A. B., M. A. Semikhatov, and I. M. Gorokhov. 2012. The Sr isotope composition of the world ocean, marginal and inland seas: Implications for the Sr isotope stratigraphy. Stratigraphy and Geological Correlation 20:501–515.

Limburg, K. E., T. A. Hayden, W. E. Pine, M. D. Yard, R. Kozdon, and J. W. Valley. 2013. Of travertine and time: Otolith chemistry and microstructure detect provenance and demography of endangered humpback chub in Grand Canyon, USA. PLoS ONE 8:e84235.

Lobo, L., R. Pereiro, and B. Fernández. 2018. Opportunities and challenges of isotopic analysis by laser ablation ICP-MS in biological studies.

Ludsin, S. A., B. J. Fryer, and J. E. Gagnon. 2006. Comparison of Solution-Based versus Laser Ablation Inductively Coupled Plasma Mass Spectrometry for Analysis of Larval Fish Otolith Microelemental Composition. Transactions of the American Fisheries Society 135:218–231.

Mallela, J., J. Hermann, R. P. Rapp, and S. M. Eggins. 2011. Fine-scale phosphorus distribution in coral skeletons: Combining X-ray mapping by electronprobe microanalysis and LA-ICP-MS. Coral Reefs 30:813–818.

Markulin, K., M. Peharda, R. Mertz-Kraus, B. R. Schöne, H. Uvanović, Ž. Kovač, and I. Janeković. 2019. Trace and minor element records in aragonitic bivalve shells as environmental proxies. Chemical Geology 507:120–133.

Martin, J., G. Bareille, S. Berail, C. Pecheyran, F. Daverat, N. Bru, H. Tabouret, and O. Donard. 2013. Spatial and temporal variations in otolith chemistry and relationships with water chemistry: A useful tool to distinguish Atlantic salmon Salmo salar parr from different natal streams. Journal of Fish Biology 82:1556–1581.

El Meknassi, S., Dera, T. Cardone, M. De Rafélis, C. Brahmi, and V. Chavagnac. 2018. Sr isotope ratios of modern carbonate shells: Good and bad news for chemostratigraphy. Geology 46:1003–1006.

Miller, J. A., B. K. Wells, S. M. Sogard, C. B. Grimes, and G. M. Cailliet. 2010, November 4. Introduction to proceedings of the 4th International Otolith Symposium. Springer.

Mokadem, F., I. J. Parkinson, E. C. Hathorne, P. Anand, J. T. Allen, and K. W. Burton. 2015. High-precision radiogenic strontium isotope measurements of the modern and glacial ocean:Limits on glacial-interglacial variations in continental weathering. Earth and Planetary Science Letters 415:111–120.

Morales-Nin, B., and A. J. Geffen. 2015. The use of calcified tissues as tools to support management: The view from the 5th international otolith symposium. Page ICES Journal of Marine Science.

Muhlfeld, C. C., S. R. Thorrold, T. E. McMahon, B. Marotz, and B. Gillanders. 2012. Estimating westslope cutthroat trout (Oncorhynchus clarkii lewisi) movements in a river network using strontium isoscapes. Canadian Journal of Fisheries and Aquatic Sciences 69:906–915.

Oschmann, W. 2009, December 23. Sclerochronology: Editorial. Springer.

Outridge, P., S. Chenery, J. Babaluk, and J. Reist. 2002. Analysis of geological Sr isotope markers in fish otoliths with subannual resolution using laser ablation-multicollector-ICP-mass spectrometry. Environmental Geology 42:891–899.

Paton, C., J. Hellstrom, B. Paul, J. Woodhead, and J. Hergt. 2011. Iolite: Freeware for the visualisation and processing of mass spectrometric data. Journal of Analytical Atomic Spectrometry 26:2508–2518.

Pozebon, D., G. L. Scheffler, and V. L. Dressler. 2017. Recent applications of laser ablation inductively coupled plasma mass spectrometry (LA-ICP-MS) for biological sample analysis: a follow-up review.

Prohaska, T., J. Irrgeher, and A. Zitek. 2016. Simultaneous multi-element and isotope ratio imaging of fish otoliths by laser ablation split stream ICP-MS/MC ICP-MS.

R Development Core Team, R. 2011. R: A Language and Environment for Statistical Computing. R Foundation for Statistical Computing.

Rodda, C., P. Mayewski, A. Kurbatov, E. Aizen, V. Aizen, E. Korotkikh, N. Takeuchi, K. Fujita, K. Kawamura, and A. Tsushima. 2019. Seasonal variability in a 1600 year-long ice core chemical record, Pamir Mountains, Central Asia. arXive arXiv:1910.

Rubenstein, D. R., and K. A. Hobson. 2004. From birds to butterflies: animal movement patterns and stable isotopes. Review TRENDS in Ecology and Evolution 19.

Rüggeberg, A., J. Fietzke, V. Liebetrau, A. Eisenhauer, W. C. Dullo, and A. Freiwald. 2008. Stable strontium isotopes (δ88/86Sr) in cold-water corals - A new proxy for reconstruction of intermediate ocean water temperatures. Earth and Planetary Science Letters 269:570–575.

Secor, D. H. 2010. Is otolith science transformative? New views on fish migration. Environmental Biology of Fishes 89:209–220.

Seeley, M., N. Miller, and B. Walther. 2015. High resolution profiles of elements in Atlantic tarpon (Megalops atlanticus) scales obtained via cross-sectioning and laser ablation ICPMS: a literature survey and novel approach for scale analyses. Environmental Biology of Fishes 98:2223–2238.

Sirot, C., F. Ferraton, J. Panfili, A.-R. Childs, F. Guilhaumon, and A. M. Darnaude. 2017. elementr: An R package for reducing elemental data from LA-ICPMS analysis of biological calcified structures. Methods in Ecology and Evolution 8:1659–1667.

Sturrock, A. M., E. Hunter, J. A. Milton, R. C. Johnson, C. P. Waring, and C. N. Trueman. 2015a. Quantifying physiological influences on otolith microchemistry. Methods in Ecology and Evolution 6:806–816.

Sturrock, A. M., J. D. Wikert, T. Heyne, C. Mesick, A. E. Hubbard, T. M. Hinkelman, P. K. Weber, G. E. Whitman, J. J. Glessner, and R. C. Johnson. 2015b. Reconstructing the migratory behavior and long-term survivorship of juvenile Chinook salmon under contrasting hydrologic regimes. PLoS ONE 10:e0122380.

Thornton, E. K. 2011. Reconstructing ancient Maya animal trade through strontium isotope (87Sr/86Sr) analysis. Journal of Archaeological Science 38:3254–3263.

Volk, E. C., A. Blakley, S. L. Schroder, and S. M. Kuehner. 2000. Otolith chemistry reflects migratory characteristics of Pacific salmonids: using otolith core chemistry to distinguish maternal associations with sea and freshwaters. Fisheries Research 46:251–266.

Walther, B. D. 2019. The art of otolith chemistry: Interpreting patterns by integrating perspectives. Pages 1643–1658 Marine and Freshwater Research. CSIRO.

Walther, B. D., and S. R. Thorrold. 2008. Continental-scale variation in otolith geochemistry of juvenile American shad (Alosa sapidissima). Canadian Journal of Fisheries and Aquatic Sciences 65:2623–2635.

Walther, B. D., S. R. Thorrold, and J. E. Olney. 2008. Geochemical Signatures in Otoliths Record Natal Origins of American Shad. Transactions of the American Fisheries Society 137:57–69.

Wang, C.-H., B. D. Walther, and B. M. Gillanders. 2019. Introduction to the 6th International Otolith Symposium. Marine and Freshwater Research 70:i.

Warter, V., and W. Müller. 2017. Daily growth and tidal rhythms in Miocene and modern giant clams revealed via ultra-high resolution LA-ICPMS analysis — A novel methodological approach towards improved sclerochemistry. Palaeogeography, Palaeoclimatology, Palaeoecology 465:362–375.

Wassenaar, L. I., and K. A. Hobson. 1998. Natal origins of migratory monarch butterflies at wintering colonies in Mexico: new isotopic evidence. Proceedings of the National Academy of Sciences of the United States of America 95:15436–15439.

Weber, M., F. Lugli, K. P. Jochum, A. Cipriani, and D. Scholz. 2018. Calcium Carbonate and Phosphate Reference Materials for Monitoring Bulk and Microanalytical Determination of Sr Isotopes. Geostandards and Geoanalytical Research 42:77–89.

Webster, M. S., P. P. Marra, S. M. Haig, S. Bensch, and R. T. Holmes. 2002. Links between worlds: unraveling migratory connectivity. Trends in Ecology & Evolution 17:76–83.

Willmes, M., K. M. Ransom, L. S. Lewis, C. T. Denney, J. J. G. Glessner, and J. A. Hobbs. 2018. IsoFishR: An application for reproducible data reduction and analysis of strontium isotope ratios (87 Sr/ 86 Sr) obtained via laser-ablation MC-ICP-MS. PLoS ONE 13.

Woodcock, S. H., C. A. Grieshaber, and B. D. Walther. 2013. Dietary transfer of enriched stable isotopes to mark otoliths, fin rays, and scales. Canadian Journal of Fisheries and Aquatic Sciences 70:1–4.

Woodhead, J., S. Swearer, J. Hergt, and R. Maas. 2005. In situ Sr-isotope analysis of carbonates by LA-MC-ICP-MS: interference corrections, high spatial resolution and an example from otolith studies. Journal of Analytical Atomic Spectrometry 20:22–27.

Zhou, H., Y. xing Feng, J. xin Zhao, C. C. Shen, C. F. You, and Y. Lin. 2009. Deglacial variations of Sr and 87Sr/86Sr ratio recorded by a stalagmite from Central China and their association with past climate and environment. Chemical Geology 268:233–247.

Zitek, A., M. Sturm, H. Waidbacher, and T. Prohaska. 2010. Discrimination of wild and hatchery trout by natural chronological patterns of elements and isotopes in otoliths using LA-ICP-MS. Fisheries Management and Ecology 17:435–445.

